# Dystrophic microglia are a disease associated microglia morphology in the human brain

**DOI:** 10.1101/2020.07.30.228999

**Authors:** Ryan K. Shahidehpour, Rebecca E. Higdon, Nicole G. Crawford, Janna H. Neltner, Eseosa T. Ighodaro, Ela Patel, Douglas Price, Peter T. Nelson, Adam Bachstetter

## Abstract

Microglia activation—typically described in terms of hypertrophic appearance—is a well-established feature of aging. Recent studies have suggested that microglia dystrophy, not activation, may increase the propagation of progressive neurodegenerative diseases such as Alzheimer’s disease (AD). Yet, a clear understanding of cause and consequences of dystrophic microglia is lacking. Although frequently observed in diseased brains, the appearance of dystrophic microglia in the hippocampus of individuals free of cognitive impairment suggests that microglia may be undergoing senescence with age, leading to dystrophy. Therefore, we hypothesized that chronological age could be a significant contributor to the presence of dystrophic microglia. To investigate this relationship, we employed stereological counts of total microglia, hypertrophic, and dystrophic microglia across the decades of the human lifespan. The microglia counts were performed in the frontal cortex gray and white matter and the CA1 subregion of the hippocampus in individuals without known neurodegenerative disease. There was age-associated increase in the number of dystrophic microglia in the CA1 region and the frontal cortex gray matter related to age. However, the increase in dystrophic microglia was proportional to the age-related increase in the total number of microglia, suggesting that aging alone was not sufficient to explain the presence of dystrophic microglia. We next tested if dystrophic microglia could be a disease-associated microglia phenotype. Compared to controls without neuropathology, the number of dystrophic microglia was significantly greater in aged-matched cases with either AD, dementia with Lewy bodies, or LATE-NC. These results provide evidence that healthy aging is only associated with a modest increase in dystrophic microglia and suggest that dystrophic microglia may be a disease-associated microglia phenotype. Finally, we found strong evidence for iron homeostasis changes, with increases in ferritin light chain, in dystrophic microglia compared to ramified or hypertrophic microglia. Based on our findings, microglia dystrophy, and not hypertrophic microglia, are the disease-associated microglia morphology. Future work is required to understand the links between the increase in dystrophic microglia and neurodegenerative disorders.

## Introduction

Inflammation and cellular senescence are hallmarks of aging [1, 2]. Almost two decades ago, dystrophic microglia were described, with beading and fragmentation of the branches of the microglia [3]. While the cellular processes appear to be fragmented, they are actually intact, with the bead-like portions connected by thin (0.18 μm) channels [4]. In contrast to the hypertrophic microglia often seen following CNS injury, the dystrophic microglia were proposed to be a form of microglia senescence [3].

While there is no single specific marker of cellular senescence, a handful of markers, such as p16^INK4a^, and p21^WAF1/Cip1^, have some affinity for identifying senescent cells [5]. Using a p16^INK4a^ approach to target the removal of senescent cells in a mouse model of tauopathy resulted in reduced tau pathology, neuronal degeneration, and cognitive deficits [6]. Given the necessary cellular stressors, a microglia can become senescent/dystrophic, as recently reported following a TBI in aged mice, where an increase in the senescent markers p16^INK4a^ and p21^WAF1/Cip1^ were seen in microglia [7].

Throughout the body, cellular senescence is associated with the secretion of inflammatory mediators, defined as the senescence-associated secretory phenotype (SASP). The SASP includes the production of matrix metalloproteinases, interleukins, chemokines, nitric oxide, and ROS [5]. Intriguingly, these findings suggest that the senescent/dystrophic microglia, and not the hypertrophic microglia, could produce the chronic inflammatory mediators associated with neuroinflammation and inflammaging. Even a small number of senescent cells in any organ can contribute to disease and by the spread of the senescence phenotype to neighboring healthy cells [8].

The hypothesis that dystrophic microglia is an age-associated microglia phenotype has not been experimentally tested. While cellular senescence generally increases with age, it can occur at any stage of life in response to stressors [5]. This led our first question: are dystrophic microglia associated with chronological age in people? We hypothesized that with increasing years, there would be an increasing proportion of dystrophic microglia. Previous work, including our own, has found dystrophic microglia in aged humans without neurodegenerative pathology [3, 9, 10].

In contrast to the view that dystrophic microglia are purely an age-related change in microglial phenotype, there is compelling evidence that dystrophic microglia are more closely associated with neurodegenerative disease. Previous studies identified dystrophic microglia in people with age-related neurodegenerative disease, including Alzheimer’s disease (AD) [4, 9–12], Down syndrome [11, 13], Huntington disease [14], dementia with Lewy bodies [9, 15], limbic-predominant age-related TDP-43 encephalopathy (LATE) [9], and multiple sclerosis [16]. These findings lead to our second question: is increased dystrophic microglia a disease associated phenomenon? We hypothesized that the absolute numbers, and/or percentage of dystrophic microglia, would be greater in people with neurodegenerative disease than age-matched controls.

To address these questions, we studied brains from the University of Kentucky Depts of Pathology and the UK-ADRC biobank, covered the adult lifespan from 10-90+ years of age. Stereological counts of the total number of microglia, number of hypertrophic microglia, and the number of dystrophic microglia were conducted in three brain regions: hippocampal CA1, frontal cortex gray matter, and white matter. We found that dystrophic microglia were increasing with age in the hippocampus and frontal cortex. However, with neurodegenerative pathology, the percentage of microglia observed to be dystrophic was much greater than that seen with aging. These results suggest that aging in the absence of neurodegenerative disease is only associated with modest increases in dystrophic microglia. In contrast we found a large increase in dystrophic microglia in brains with neurodegenerative pathology compared to control brains. Previous studies have suggested dysfunctional iron metabolism could lead to increased oxidative stress contributing to the degeneration of microglia [17]. In brains with neurodegenerative pathology, we found a strong association of ferritin light chain (FTL) with dystrophic microglia compared to other microglia phenotypes. FTL is the major protein responsible for storing intercellular iron. These findings highlight the iron metabolic pathway as a potential molecular mechanism that leads to the dystrophic disease associated with microglia morphology.

## Materials and Methods

### Human subjects

Tissue samples that contained the hippocampus or frontal cortex were acquired from the University of Kentucky biobank, and from the University of Kentucky Department of Pathology and Laboratory Medicine. The reason for using these latter cases was to incorporate data from younger subjects. Cases were selected by the investigators (JHN and PTN), with the exclusion criteria of pathologically confirmed neurodegenerative disease: specifically, but not limited to, advanced disease pathology associated with Alzheimer’s disease, dementia with Lewy bodies, LATE, and vascular dementia. Demographic data are presented in **Table 1**. The matching hippocampus and frontal cortex were not available for all cases. Three additional cases of LATE neuropathological change (LATE-NC), not part of the original cohort where used for the IBA1/FTL analysis.

**Table 1:**
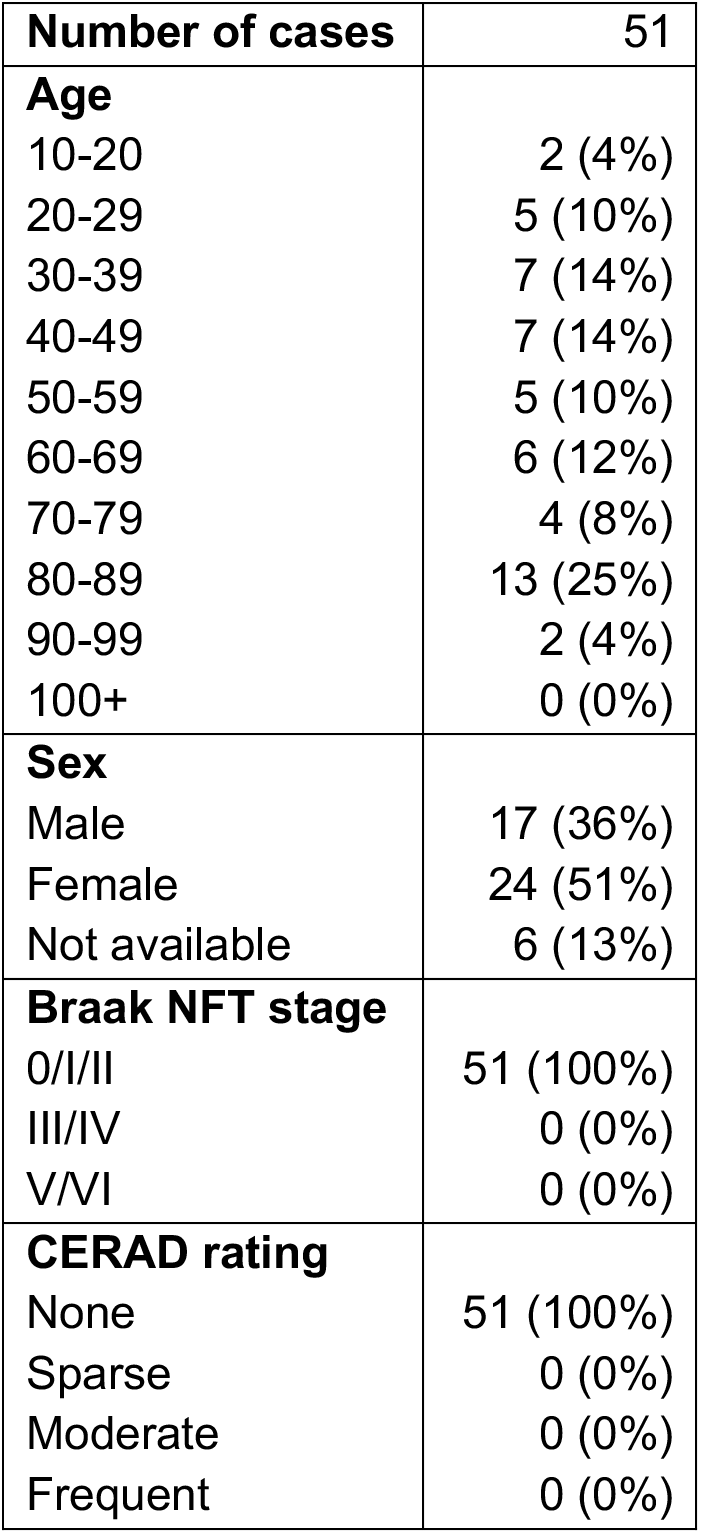
Aging case series characteristics.

| Number of cases | 51 |
| --- | --- |
| **Age** |  |
| 10-20 | 2 (4%) |
| 20-29 | 5 (10%) |
| 30-39 | 7 (14%) |
| 40-49 | 7 (14%) |
| 50-59 | 5 (10%) |
| 60-69 | 6 (12%) |
| 70-79 | 4 (8%) |
| 80-89 | 13 (25%) |
| 90-99 | 2 (4%) |
| 100+ | 0 (0%) |
| **Sex** |  |
| Male | 17 (36%) |
| Female | 24 (51%) |
| Not available | 6 (13%) |
| **Braak NFT stage** |  |
| 0/I/II | 51 (100%) |
| III/IV | 0 (0%) |
| V/VI | 0 (0%) |
| **CERAD rating** |  |
| None | 51 (100%) |
| Sparse | 0 (0%) |
| Moderate | 0 (0%) |
| Frequent | 0 (0%) |

### Immunostaining

Immunohistochemical (IHC) staining for IBA1 was completed as previously described [9]. Briefly, microwave antigen retrieval (6 min (power 8) using citrate buffer (Declare buffer, Cell Marque; Rocklin, CA)) was done following de-paraffinization on 10-μm-thick tissue sections. Endogenous peroxidases were quenched in 3% H_2_O_2_ in methanol for 30 min. Sections were blocked in 5% normal goat serum at room temperature for 1 hour. Sections were incubated in primary antibodies IBA1 (rabbit polyclonal, 1:1,000, Wako Catalog no. 019-19741, AB_839504); 20-24 hours at 4°C. A biotinylated secondary antibody (Vector Laboratories) was amplified using avidin-biotin substrate (ABC solution, Vector Laboratories catalog no. PK-6100), followed by color development in Nova Red, or DAB (Vector Laboratories). The double-label immunofluorescence was completed as previously described [18]. Briefly, microwave antigen retrieval (6 min (power 8) using citrate buffer (Declare buffer, Cell Marque; Rocklin, CA)) was done following de-paraffinization on 8-μm-thick tissue sections. Sections were incubated for 45 sec at room temperature in a 1x solution of TrueBlack (Cat. #23007, Biotium, Fremont, CA) prepared in 70% ethanol, to reduce auto-fluorescence. Following blocking in 5% normal goat serum, the sections were incubated in IBA1 (rabbit polyclonal, 1:1,000, Wako Catalog no. 019-19741, AB_839504), and FTL (mouse clone D9, 1:100, Santa Cruz, Catalog no sc-74513, AB_1122837) for 20-24 hours at 4°C. Sections were incubated in secondary antibodies conjugated to Alexa Fluor probes (Life Technologies; Carlsbad, CA) diluted 1:200 were room temperature for 1 hour. Control sections were included for each case with that omitted one or both of the primary antibodies.

### Quantitative image analysis

Briefly, the Zeiss Axio Scan Z.1 digital slidescanner was used to image the entire stained slide at 40x magnification to create a single high-resolution digital image. Halo software (version 2.3; Indica labs) was used to view the images. Following the fractionator method of stereology, we used the Halo software to generate counting frames 250 × 250μm, with a 150μm gap between counting frames using systematic random sampling. A total of 10-20 counting frames were quantified to estimate the number of microglia per unit area in the three brain regions. Classification of microglia as either hypertrophic or dystrophic followed our previously described criteria [9]. Example photomicrographs of representative cells defined as either hypertrophic or dystrophic microglia is shown in **Figure 1**. Results were confirmed by two independent observers (REH, and NGC) blind to experimental conditions. Data presented is from REH’s quantification. Numbers of dystrophic microglia in cases with neurodegenerative disease were generated as part of our prior published study [9], but follow methods described above.

**Figure 1:**
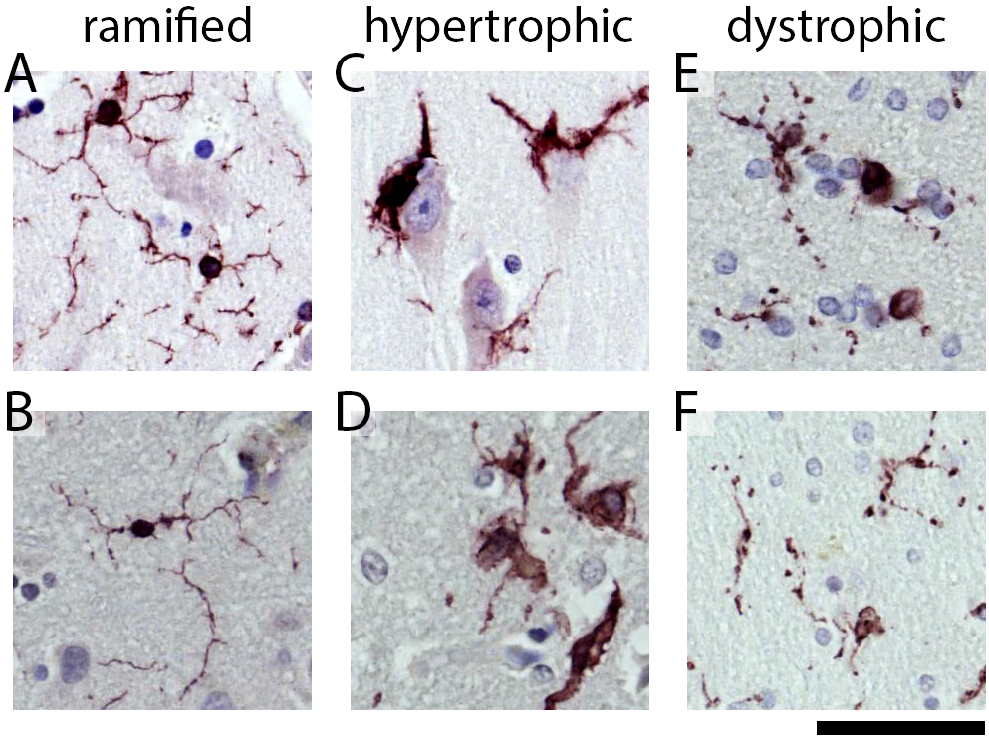
Representative examples of the microglia morphological appearances seen in the cases. **(A, B)** Shows examples of the ramified microglia. Hypertrophic microglia are shown in **(C, D)**. In **(C)**, microglia can be seen surrounding the soma of a pyramidal neuron. **(E)** shows dystrophic microglia with swellings, beaded, and discontinuous processes. **(F)** shows an example of dystrophic microglia that are small and de-ramified, with beaded processes. The scale bar is 50 μm.

For the IBA1 and FTL colocalization analysis a Zeiss Axio Scan Z.1 digital slidescanner was used to image the entire stained slide at 40x magnification to create a single high-resolution digital image. Halo software (version 2.3; Indica labs) was used to view the images. Using HALO imaging software and viewing only the IBA1 channel when counting and determining the morphology of the cells, we generated an ROI around each microglia, and define it as ramified, hypertrophic or dystrophic. We identified at random 10 microglia from each morphology for each of three cases of LATE-NC. We then used the area colocalization FL algorithm (Halo software, version 2.3; Indica labs) on the ROIs to determine the area of colocalization of IBA1 and FTL for the three microglia morphologies.

### Statistics

JMP Pro software version 14.0 (SAS institute, Cary, NC, USA) or GraphPad Prism software version 8.0 was used to generate graphs and for statistical analysis. Linear regression and Spearman r were used to compare the effect of age on microglia morphological state. Mean ± 95% confidence interval are shown for the regressions, as well the data point for each case. Standard least squares model was used to compare hypertrophic and dystrophic microglia by was corrected for age.

## Results

Microglia can be classified into distinct morphological phenotypes using IBA1 immunohistochemistry. As previously described [9], the morphological phenotypes include ramified microglia, which are thought to be a healthy, or homeostatic, phenotype **(Fig 1 A, B)**. In contrast, the hypertrophic microglia morphology has historically been associated with a reactive microglia response. This phenotype is typically observed following acute neuronal injury and surrounding amyloid plaques **(Fig 1 C, D)**. As previously described, dystrophic microglia morphology refers to a number of morphological changes affecting the cytoplasmic processes such as spheroidal swelling, de-ramified, beaded, discontinuous, or tortuous processes **(Fig 1 E, F)** [2,3,4]. A trained scientist can quantify these three distinct microglia morphologies. Thus far, we have been unsuccessful in training computer algorithms, including neural network algorithms, in detecting these phenotypes accurately. Therefore, we adopted a design-based stereological approach and replicated the quantification using two observers blind to the experimental conditions (R.E.H and N.G.C).

### Age affects microglial morphology in hippocampal subregions of the human brain

To better understand the link between dystrophic microglia and age, we began examining the CA1 region of the hippocampus. Regardless of microglia morphology, we found a strong correlation for an increase in the number of microglia with greater age **(Fig 2A)**. Similarly, we also saw a rise in the number of hypertrophic microglia **(Fig 2B)** and dystrophic microglia **(Fig 2C)** with age. We next tested if there was a change in the proportion of microglia that were hypertrophic or dystrophic as a function of age. We found that when the hypertrophic microglia were compared as a percentage of the total microglia there was a strong positive correlation for an increased percentage of hypertrophic microglia with age **(Fig 2D)**. We did not find evidence in support of our hypothesis that dystrophic microglia were associated with age, as there was a lack of a correlation for the percentage of dystrophic microglia to total microglia with age **(Fig 2E)**. That is, while the total number of dystrophic microglia increases with age in the CA1 region of the hippocampus **(Fig 2C)**, this change can be accounted for by the age-related increase in the total number of microglia **(Fig 2A)**. The lack of an association with age and dystrophic microglia is in contrast to the hypertrophic microglia, which are increasing with age **(Fig 2D)**.

**Figure 2:**
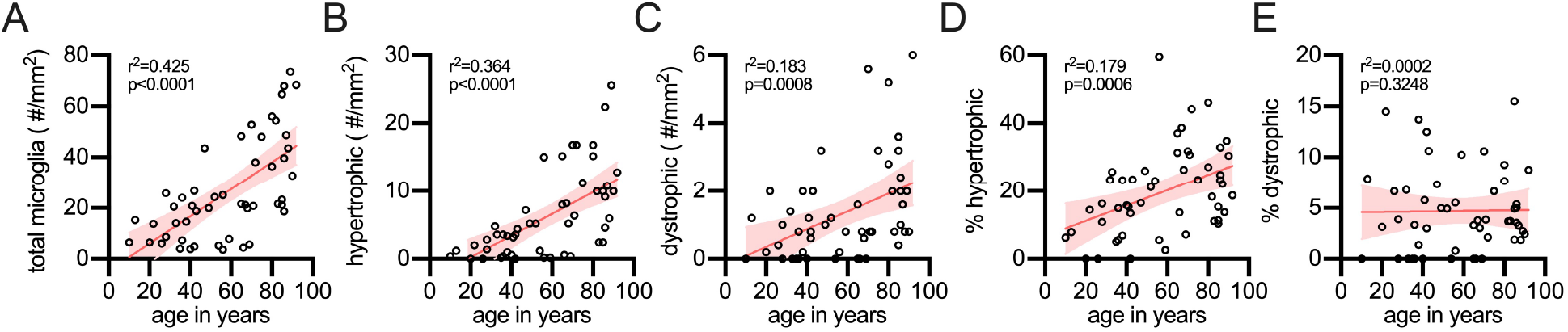
Effect of age on microglial morphology in the CA1 region of the hippocampus. **(A)** The total number of microglia, regardless of morphology, were found to increase with age in the hippocampus. The total number of hypertrophic **(B)**, as well as dystrophic microglia **(C)**, were also found to increase in the hippocampus with age. The percentage of the total microglia which are hypertrophic was found to increase with age **(D)**, while the increase in percentage of microglia that were dystrophic (**E**) was proportional to the total number of microglia, and thus did not increase with age. Each circle is a unique person.

### Changes in microglial morphology in neocortical gray matter regions of the brain

Next, we moved from the limbic system to evaluate changes in the spatial distribution of dystrophic microglia in the neocortex. Investigation of microglia in frontal lobe gray matter demonstrated several striking differences in comparison to the hippocampus. While the total number of microglia was found to increase with age **(Fig 3A)**, there was no age-related increase in the number of hypertrophic microglia in the frontal cortex **(Fig 3B)**. Counts of the number of dystrophic microglia were correlated with age **(Fig 3C)**. As a proportion of the total microglia, hypertrophic microglia did not increase with age **(Fig 3D)**, while there was an increase in the percentage of dystrophic microglia with age **(Fig 3E)**.

**Figure 3:**
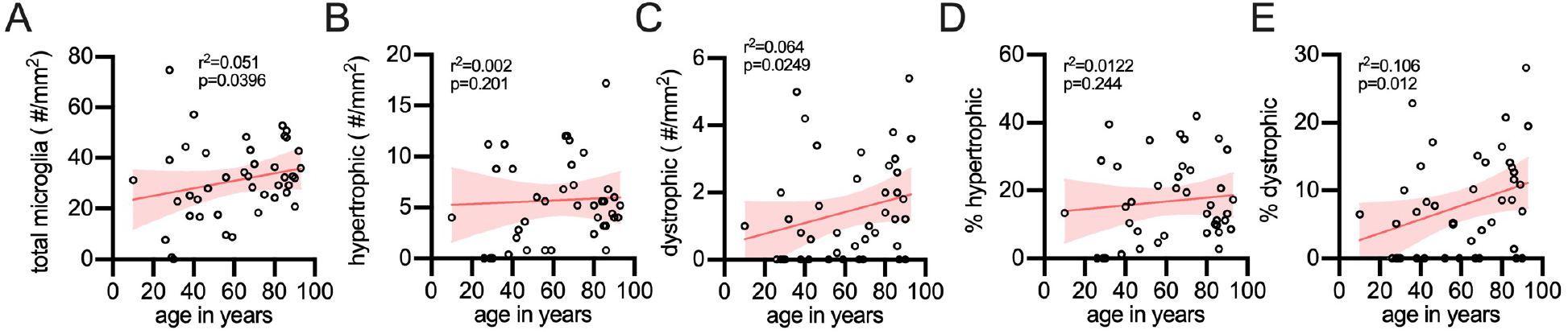
Stereological quantification of microglia morphology in the frontal cortex gray matter as a function of age. **(A)** The total number of microglia were found to increase with age. **(B)** However, hypertrophic microglia showed no age-related increase. **(C)** Dystrophic microglia were found to increase with age. **(D)** The percentage of total microglia that were hypertrophic did not increase with age; whereas, **(E)** dystrophic did increase with age in the frontal cortex gray matter. Each circle is a unique person.

### Changes in microglial morphology in white matter regions of the brain

In addition to the limbic and neocortical gray matter, we also quantified microglia in the white matter of the frontal cortex. We found that the total number of microglia **(Fig 4A)**, the number of hypertrophic microglia **(Fig 4B)**, and the number of dystrophic microglia **(Fig 4C)** did not increase with age in the white matter. Also, the percentage of hypertrophic microglia **(Fig 4D)**, or dystrophic cells **(Fig 4E)** were unchanged with age.

**Figure 4:**
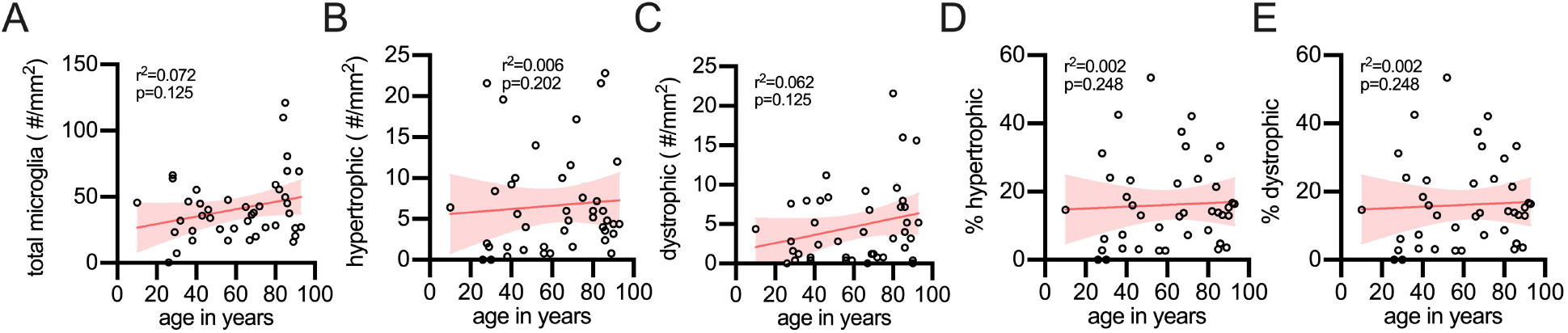
Effect of age on microglial morphology in the frontal cortex white matter. The number of total microglia **(A)**, as well as the number of hypertrophic **(B)** and dystrophic **(C)** microglia did not increase in the white matter of the frontal cortex. Similarly, there was no change in the percentage of hypertrophic **(D)** or dystrophic microglia **(E)**.

### Dystrophic microglia are a disease-associated microglia morphology

Given that we found little evidence of dystrophic microglia increasing with age, we next asked if dystrophic microglia could be characteristic of a disease-associated microglia morphology. To address this hypothesis we compared changes in dystrophic microglia in the hippocampal CA1 region from people that were greater than 65+ years old without neurodegenerative pathology, to 65+ years old people with either Alzheimer’s disease (AD), limbic-predominant age-related TDP-43 encephalopathy (LATE), or dementia with Lewy bodies (DLB) from a preexisting data set [9] **(Table 2)**. Strikingly we found that hypertrophic microglia, as a percentage of the total microglia population, were not increased in the individuals with neurodegenerative disease compared to controls **(Fig 5A)**. Instead, we found that the percentage of microglia defined as hypertrophic decreased in the CA1 region of the hippocampus in people with neurodegenerative pathology compared to those without neurodegenerative pathology. These results are in contrast to what was found for dystrophic microglia. In those cases, with neurodegenerative pathology, on average, 45% of the microglia were found to be dystrophic. This is in comparison to the cases without neurodegenerative pathology, where on average, only 9% of the microglia were dystrophic **(Fig 5A)**. While for the person in the neurodegenerative group level, the range dystrophic microglia were broad (0-100%), there was no statistical difference between the different neurodegenerative disease. These results suggest that the dystrophic microglia phenotype represents a disease-associated microglia morphology.

**Table 2:** Neurodegenerative series characteristics.

| Table 2: Neurodegenerative series characteristics |  |  |  |  |
| --- | --- | --- | --- | --- |
| Experimental group | Number of cases | Sex |  | Age range |
|  |  | female | male |  |
| healthy control | 34 | 16 | 13 | 65-93 |
| AD | 8 | 3 | 5 | 65-85 |
| DLB | 11 | 2 | 9 | 65-97 |
| LATE | 9 | 5 | 4 | 65-93 |

**Figure 5:**
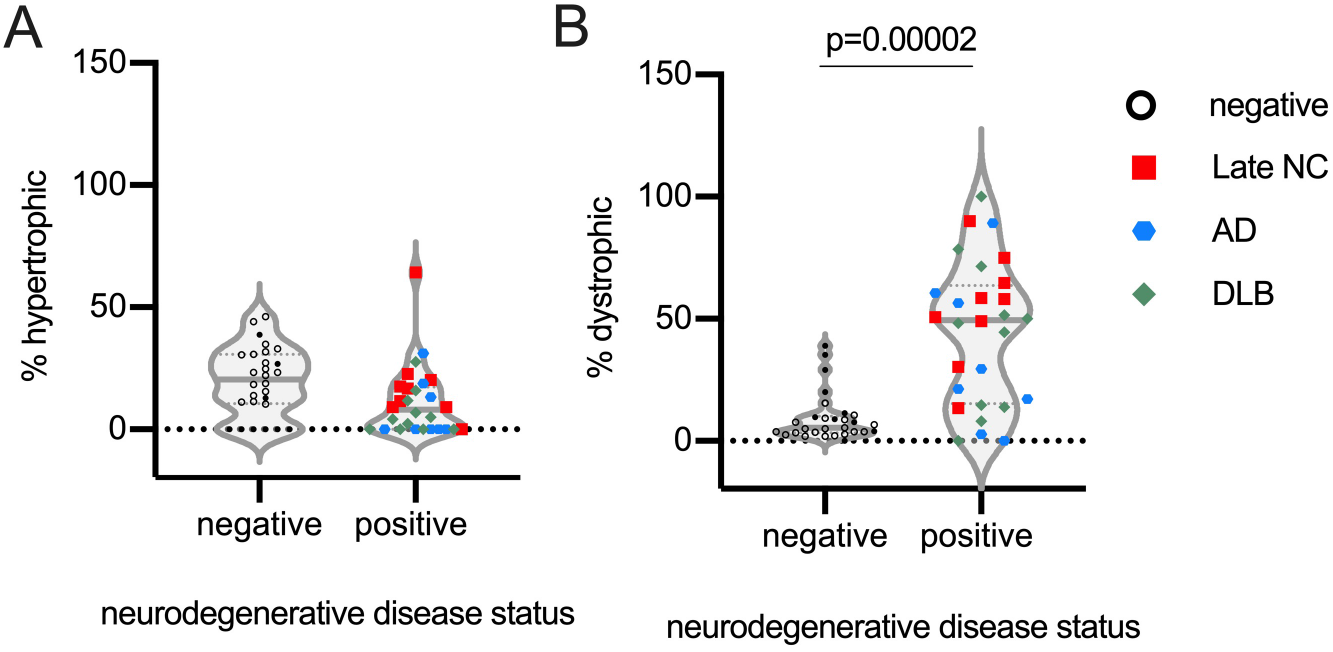
Association of dystrophic microglia with neurodegenerative disease. Comparing the percentage of dystrophic microglia in the CA1 region in the 65+ year old cases free of advanced neurodegenerative disease pathology (negative) to those cases with advanced stage neurodegenerative disease pathology (positive) **(A)** showed a significant decrease in the number hypertrophic microglia (p=0.0998), and a **(B)** significant increase in dystrophic microglia (p=0.00002) in the cases with advanced stage neurodegenerative disease. Open symbols are from cases from the current study. Closed symbols are from our prior study [9].

### Iron metabolism as a molecular mechanism for the dystrophic disease associated microglia morphology

Single-cell and single-nuclei transcriptomic approaches have defined a gene signature in microglia associated with neurodegenerative pathology [19–23]. Exploring these datasets, we identified genes associated with the iron homeostasis enriched in aging and disease-associated microglia. While several gene members of the iron homeostatic pathway were differentially expressed in the aging and disease associated microglia, FTL was one of the iron pathway genes found to be increased across multiple datasets. In addition, prior studies have observed dystrophic microglia labeled by FTL [10, 16]. Therefore, we next hypothesized that dystrophic microglia in the aged brain with neurodegenerative disease pathology would be FTL positive. To test this, we used three LATE-NC cases and double labeled those cases using IBA1 and FTL **(Fig 6A)**. Using the IBA1 channel alone, we identified 10 ramified microglia, 10 hypertrophic microglia, and 10 dystrophic microglia per case. Using HALO imaging software, we determined the area of colocalization of IBA1 and FTL for each microglia **(Fig 6B)**. In support of our approach, we found that the intensity of the IBA1 staining in the cells classified as hypertrophic was greater than the cells classified as ramified or dystrophic microglia **(Fig 6C;** p<0.0001 Tukey’s test). We then asked, do the cells we defined as dystrophic have greater amounts of FTL. Comparing the area of IBA1+FTL+ colocalization, we found that the dystrophic microglia had much more FTL as a proportion of the overall cell than the ramified or hypertrophic microglia **(Fig 6D** p<0.0001 Tukey’s test).

**Figure 6:**
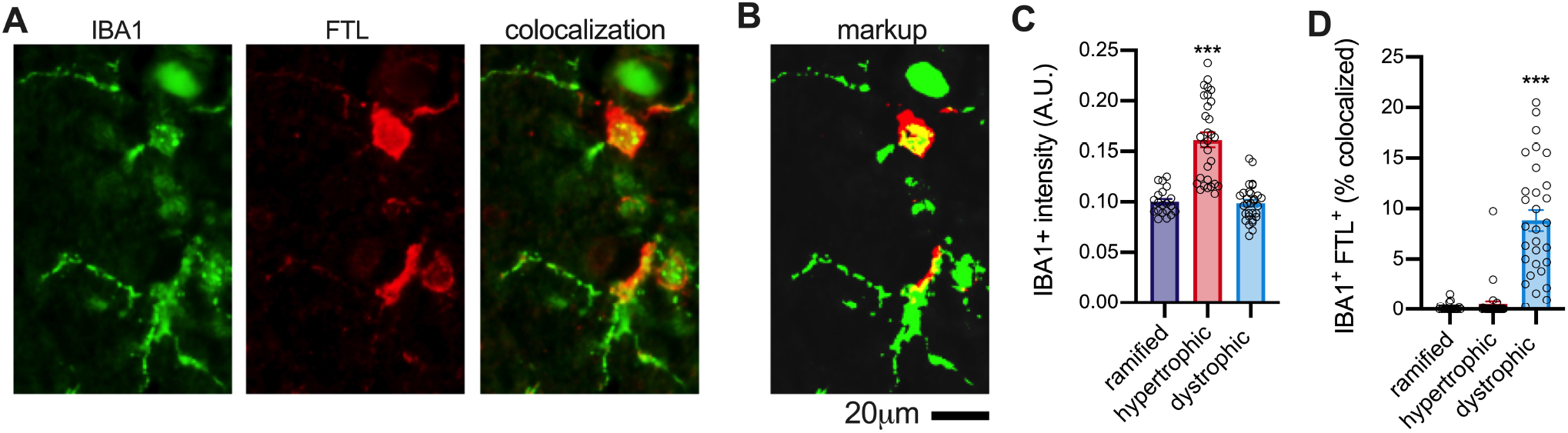
Dystrophic microglia have a strong tendency to be double-labeled for FTL in the hippocampus of aged humans. A quantitative cellular localization of FTL found colocalization FTL with microglia. FTL did not appear to be staining all microglia; therefore, we sought to see if there was any association of FTL with the different microglia phenotypes we recently characterized. (**A**) Shows an example of the IBA1^+^FTL^+^ staining. (**B**) Shows the HALO generated markup, where green is IBA1, red is FTL, and yellow is the area where the two proteins are colocalized. (**C**) Shows that there was greater IBA1 staining intensity in the hypertrophic microglia compared to the two different morphologies (p<0.0001, Tukey Test). (**D**) A high degree of IBA1^+^FTL^+^ area of colocalization in the dystrophic microglia compared to the ramified or hypertrophic morphologies (p<0.0001, Tukey Test). Circles are individual microglia, from 3 different aged brains.)

## Discussion

In the nearly two decades since dystrophic microglia were first described [3], the causes and consequences of this unique cellular morphology remains undefined (for review see: [17]). Like other types of cellular senescence that occur with age, we asked if aging was sufficient to drive microglia towards a dystrophic microglia morphology in the absence of disease. To begin to address these critical knowledge gaps, we used a series of autopsy cases covering the human lifespan to define the association of dystrophic microglia with age. We found only limited evidence that dystrophic microglia increased with age in the absence of neurodegenerative pathology. We saw in the hippocampal CA1 region a robust age-related increase in the total number of microglia, as well as the total number of hypertrophic and dystrophic microglia. However, as a percentage of the total number of microglia, only the hypertrophic microglia morphology was found to increase with age. At the same time, there was no relationship between age and the percentage of dystrophic microglia. Similar trends were found in the gray matter of the neocortex. After observing a general lack of an association with age and dystrophic microglia, we next asked if dystrophic microglia are a disease associated microglia morphology. In contrast to the aged brain without neurodegenerative disease, we found strong evidence that dystrophic microglia are a disease associated microglia morphology. While there is likely more than one mechanism that push microglia into a dystrophic phenotype, we identified iron metabolism as a potential molecular mechanism warranting future study.

We used a two-dimensional stereological approach to quantify the number of microglia by their morphological appearance. In the scarce tissue samples used in our study, it was not feasible to use three-dimensional stereological approaches, such as the optical fractionator, which would account for changes with tissue shrinkage caused by fixation [24, 25]. There is also subjectivity associated with our methods as the observers – who were blind to the experimental groups – needed to classify a cell as hypertrophic, dystrophic, or other. We used a number of approaches to limit the subjectivity. For example, we repeated the assay using two independent scientists. We also tested intra-observer variability by having the same scientist replicate a subset of their microglia counts. Our results are comparable to those of Davies et al., which used morphometric approaches to quantify microglia arborization and microglia coverage and found a decrease in microglia complexity and area of the brain covered by microglia in the AD brain compared to the matching control brains [26]. Also, using CD68 and MHCII it was shown that there was an age-related decrease in these markers in AD brains, while there was no change in the control cases with age [27], which is in agreement with our results. While we acknowledge the limitations associated with our approach, our study provides a quantitative assessment of microglia number by morphology across the human lifespan, which has not been previously reported.

Our results that dystrophic microglia are the dominant disease-associated microglia morphology may appear counter to the current assumptions that neurodegenerative disease is associated with microglia activation. Indeed, as thoroughly reviewed by Hopperton et al., the vast majority of studies do find that microglia/macrophages are increased in AD brains compared to control [28]. However, the likelihood that the study will report an increase in microglia/macrophages AD brains compared to control is greater when activation marker (e.g., IL-1α) or a marker of macrophages (e.g. CD163) is used [28]. Our results are not in disagreement with these findings. We believe that there may be a loss of homeostatic microglia in the neurodegenerative disease brain, with a compensatory increase in monocytes/macrophages.

We did find with age in the current study, and with neurodegenerative disease in our previous study [9] the total number of IBA1+ cells in the CA1 region of the hippocampus was increased in the brain with neurodegenerative pathology compared to the control brains. The increase in total number of microglia/macrophages may be compensatory for the loss of homeostatic coverage of the brain parenchyma by microglia. Studies using depletion approaches to eliminate microglia in animal models have clearly shown that the brain will quickly repopulate microglia to maintain a tiling of microglia across the brain [29]. However, the rate of repopulation slows following multiple rounds of depletion [29].

Comparative studies of dystrophic microglia provide important insights into the biology of this cellular morphology. In marmosets, it was found that the number of dystrophic microglia increased with age in both the limbic cortex and neocortex. However, they found this increase peaked in late middle age marmosets and declines in the oldest aged marmosets [30]. These results are in contrast to our current study, where we saw the total number of dystrophic microglia increase linearly with age. In aged chimpanzees, dystrophic microglia were found primarily in the neocortical gray matter of layer II-III [31]. While amyloid and tau neuropathological changes were found in a subset of the aged chimpanzees used in the study, quantification of dystrophic microglia was not completed, so it remains to be determined if there is an association between dystrophic microglia and amyloid and tau in the chimpanzees [31]. In *Cynomolgus macaques* exposed to chronic manganese to model Parkinson’s disease-related pathology, dystrophic microglia were found to have intracellular ferric iron [32], which is in agreement with our findings that FTL is highly expressed in dystrophic microglia. Dystrophic microglia have been shown to increase with age in tree shrews (*Tupaia belangeri)* [33], which is in agreement with our findings. Interestingly, ferritin labeled microglia was high in oxidative stress markers and had internalized hyperphosphorylated tau [33]. In mice dystrophic microglia are seldom reported; however, a recent study did find in a mouse model of P301S tauopathy, a decrease in microglia complexity in the tau mouse with age, which could be contributed to a dystrophic microglia morphology [34].

Studies of dystrophic microglia and their role in disease are faced with several limiting factors that have slowed progress in the field. One of the most prominent limitations is the lack of distinct cell markers delineating dystrophic microglia. For this reason, futures studies must attempt to find novel cellular markers of dystrophic microglia, as well as identifying specific changes in cells that contribute to disease progression. Our study was not the first to identify FTL as a marker of dystrophic microglia. Numerous previous studies have reported that microglia increase FTL in neurodegenerative disease [17, 28]. However, our work, along with Lopes et al., [10] has demonstrated that the increase in FTL may be more than a marker of a reactive microglia phenotype and could suggest a mechanism in iron dyshomeostasis that is leading to the degeneration of microglia. Single-nuclei and single-cell data sets support the hypothesis that iron pathway is differentially expressed in disease-associated microglia [19–21, 22]. Our quantitative methods to measure FTL in LATE-NC cases found that a greater amount of the dystrophic microglia was labeled with FTL in comparison to hypertrophic microglia. Recent work in aged marmosets found that FTL labeled both an activated and dystrophic phenotype [30]. Our work is not in disagreement, as we, too, can identify FTL labeled hypertrophic microglia. We think these hypertrophic microglia could represent a point of inflection in the course of disease pathology, when microglia lose their ability to safely maintain iron homeostasis and become dystrophic. Alternatively, we previously showed that FTL was found to be colocalized with TDP-43 and Tau inclusions bodies [18]. The increase in FTL could be part of a proteinopathy – including TDP-43 and Tau – leading to the degeneration of microglia and other neuronal cells. More work is needed to directly measure iron burden in FTL positive microglia from brains with and without neurodegenerative pathology. For instance, Fe(II) and Fe(III) could be measured on FACS sorted microglia from the postmortem tissue using capillary electrophoresis coupled-to-inductively-coupled plasma mass spectrometry (CE-ICP-MS)[35].

Recent studies have shown alterations in microglial morphology in the cortical white matter of subjects with neurodegeneration related to age [28]. However, the relation between dystrophic microglia and white matter degradation is still unknown. It has traditionally been held that morphological changes in white matter microglia is related to neuroinflammation characterized by an increased expression of MHCII and complement receptor C3bi [28]. Conversely, more recent studies posit that the breakdown of myelin associated with age places an increased burden on white matter microglia and that observed increases may be related to cell senescence rather than inflammation [17]. In our experiments, we did not find any changes in microglia morphology in the white matter of frontal cortex associated with healthy aging.

## Conclusions

Although a clear relationship between increased microglia dystrophy and neurodegenerative diseases is lacking, dystrophic microglia has shown decreased neuroprotective abilities and increased secretion of pro-inflammatory molecules [17]. While the link between dystrophic microglia and disease is still unknown, the presence of dystrophic microglia in diseases with varying etiology and pathology suggests that dystrophic microglia are critically involved in the shift from normal aging to disease states. Our study lends insight into the possible correlation between age and changes in microglia morphology. To date, the role that dystrophic microglia play in normal aging and neurodegeneration is still unknown. However, based on our presented findings, the loss of function from microglia dystrophy and not microglial activation is more involved in neurodegenerative disorders. Uncovering the link between genetic predispositions, specific changes to glial microenvironment and function, and disease propagation may allow future researcher to better understand the specific role of dystrophic microglia in other progressive neurodegenerative diseases.

## Acknowledgements

We are profoundly grateful to all of the study participants who make this research possible. The corresponding author, Adam Bachstetter, PhD, had full access to all of the data in the study and takes responsibility for the integrity of the data and the accuracy of the data analysis. Research reported in this publication was supported by National Institutes of Health under award numbers P30 AG028383, R35 GM124977, R21AG066865. The content is solely the responsibility of the authors and does not represent the official views of the National Institutes of Health.

## Conflict of Interest

None

## Notes

### Competing Interest Statement

The authors have declared no competing interest.

